# The development, recognition, and learning mechanisms of an animal-like neural network

**DOI:** 10.1101/535724

**Authors:** Feng Qi, Wenchuan Wu

## Abstract

Recent advances in AI show great potentials to understand the biological brain, the thinking process, and the development of intelligence. Convergent evolution shows that networks with similar architecture could acquire similar functionalities challenged by similar tasks. Here, we aim to build an animal-like neural network by discarding neuroscience-unsupported modules, such as the convolution network (CNN), error backpropagation, and softmax classification. Instead, we adopt a biologically plausible way to construct the network, using fully connected layers, taking feedback signals as imagination stream, and directly generating symbols with a texitizer. In this paper, we use a recognition and imagination (RaI) network to study the mechanisms of the animal-like vision system. The resulting network characteristics agreed fairly well with the neurophysiological measurements in previous monkey studies. First, in the network development study, RaI also experienced several critical developmental stages characterized by specificities in neuronal types, connectivity strength, and firing patterns, from the early stage of coarse salience map recognition to the mature stage of fine structure recognition. In the recognition study, we found that RaI could maintain object invariance representation under various viewing conditions by coordinated adjustment of population neurons responses. Such concerted population responses contained untangled object identity and property information that could be accurately extracted via a simple weighted summation decoder. In the learning and forgetting study, novel structure recognition was achieved by adjusting entire synapses in low magnitude while pattern specificities of original synaptic connectivity were preserved, which guaranteed a progressive learning process without disrupting the existing functionalities. This work benefits the understanding of the human neural mechanism and the building of human-like machine intelligence.

## 1 Introduction

Species, no matter animals or human beings (or even AI), can independently develop similar features in response to similar tasks such as visual recognition and auditory processing [1]. These developments share the same natural principle of ‘system free energy minimization’ [2]. Here, we build our neural network by mimicking animal brain organization [3] and expect to acquire similar network characteristics and functionalities with similar architecture and under similar training tasks. We discarded those commonly used but neuroscience-unsupported modules. (1) The CNN though has reached human-level performance in many recognition tasks [4, 5], the animal neural system does not support such kernel scanning operation across upstream neurons, and thus the biological neuronal responses do not match that of CNN units [6, 7]. Here, we use the fully connected module instead of CNN. Although the number of parameters increases substantially, the resulting activation patterns demonstrate more biological plausible characteristics than that of CNN. Like the monkey [8], RaI uses population coding to achieve the identity-preserving recognition instead of the biologically implausible max-pooling operation. (2) The error backpropagation usually allows weights to be updated throughout multiple hierarchical layers toward an optimized objective function [9]. However, current evidence seems to regard it as biologically implausible primarily for two reasons [10]. First, there is no evidence observed in the brain supporting that the feedforward and feedback connectivity meet the requirement of perfect symmetry. Second, error backpropagation requires that the global error signals should go across multiple layers to guide local weight updates, however, neuronal synaptic plasticity depends primarily on local information (the pre- and post- synaptic neuronal activity). As suggested by neuroscience studies [11, 12], the synaptic plasticity is governed by local molecular events such as the activity-dependent glutamate N-methyl-D-aspartate receptor and calcium flow instead of by feedback signals. Indeed, the real feedback connectivity provides the top-down imagination information, which is usually ignored by AI network construction due to the solid belief of error backpropagation. (3) The softmax classification is usually used to compute the probability of each category (or word) in the repository (or vocabulary) before prediction [13, 14]. However, in reality, we never evaluate all fruit categories in mind before saying ‘it is an apple’, let alone the complicated computation of the normalization term in the softmax. In our concurrent paper [15], we used a texitizer to direct output the classification by text symbols, which is more biologically plausible than softmax operation. (4) The network architectures in AI are usually not realistic, such as the very deep neural network [16] with the residual module that enables the building of 1000+ layers network [17]. Although these newly created species could achieve greater performance in a special task, it normally goes far away from realistic intelligence. For example, we can easily figure out the interconnections between several layers of visual and auditory cortices according to brain studies [18] but it is very hard to extend from image processing to language understanding based on the 1000+ layers very deep neural network. Therefore, to achieve human-like machine intelligence, one strategy is to figure out the brain (the only proved intelligence form) neural mechanism and implement it on machines in a similar way.

Hassabis highlighted the importance of understanding the biological brain in the building of intelligent machines [10]. He mentioned how the replay mechanism of episodic memory in the hippocampus had been leveraged in the training of the deep Q network (DQN). However, to make the AlphaGo work, billions of games have to be trained to cultivate a fundamental feeling or capacity (the Q value estimation) in DQN before it can generate a proper movement [19, 20]. This usually limits DQN to one specific trained task. Unlike the Q-learning, the human being has to handle various kinds of tasks and is usually required to handle new tasks even without a single practice. To flexibly address many real-time problems, one typically first acquire the tasks or goals via language and then plan in mind before carrying out actions. Our concurrent paper [15] aims to address the problem of human-like machine thinking with a language guided imagination (LGI) mechanism. In this paper, we focus on how the imagination process is formed in a biologically plausible vision system, and how virtual brain analysis techniques [10] such as single-cell recording, neuroimaging, and lesion techniques could be used to interpret the AI system.

## 2 Experiments and Results

Experimental neuroscience discovers that the visual ventral stream in the primate brain is used for core object recognition with information transmission through a feedforward, reflexive, hierarchical neural network to rapidly disentangle the object identity manifolds [8]. Inspired by this, we built a basic recognition and imagination network (Supplementary Figure 1) to provide an analogy to the animal neural system, and expected similar characteristics could be derived in response to similar recognition challenges.

Primates have distinct development stages (i.e. baby, adolescence and adult) characterized by significant differences in neural anatomy, physiology, and cognition. RaI also experienced similar development stages (i.e. the Early, Middle, and Late stages) characterized by nine critical time points (ctp) each with unique firing patterns, synaptic connectivity patterns, and recognition performance (Figure1). In the Early stage (ctp0-4), RaI took relatively few (100k) steps, to learn to capture the salience map of objects in the retina. RaI started with randomly assigned synaptic weights (ctp0) that possessed no power for recognition though digit instances were presented in the retina. Despite there was only slight synaptic plasticity in the initial stage (ctp1), RaI started to recognize the approximate locations of those digits in the retina. The initial changes of activity occurred only in the higher layers, indicating the importance of early development of other cortices (like cortices for hearing and proprioception, etc.) and their feedback signals in shaping those V1-like synapses [21]. The ctp2 witnessed a sudden drop in the number of excitatory synapses and a rapid increase of inhibitive synapses, which was also the starting point of strengthening both excitatory and inhibitory synapses. These changes allowed RaI to identify whether the digits were presented on the left or right part of the retina, which was clearly reflected by the horizontal segmentation of positive and negative synapses (Figure 1.e). Following the same trend of synaptic alterations, RaI could identify the vertical positions and orientations of objects in ctp3, with specific cluster-like distributions gradually formed (Figure 1.e). The neurons also showed remarkable firing specificities with respect to the digit’s retinal positions. In ctp4, RaI could capture the object scale information, and this was also the starting point when RaI came to identify the digit’s morphology, namely, entering the Middle stage (ctp4-6). In the Middle stage, the most striking changes occurred in ctp5, where both excitatory and inhibitory synapses stopped strengthening and began to shrink to enhance the recognition, quite similar to the pruning mechanism in the animal brain to improve the neural network efficiency [22]. At ctp6 the pruning effect was clearly demonstrated in both activity maps with significantly sparser firing patterns and synaptic weight maps with fewer but sharper distributions. At the same time, V1-like units formed a specific visual field in the retinal region. From ctp6, RaI could basically recognize and reconstruct a correct digit, and further refinements of the structural details were developed in the Late stage (ctp6-8) on a longer time scale. The synaptic excitation-inhibition ratio continually declined at a moderate speed till ctp7 where the network’s recognition performance reached a peak level. And the subsequent synaptic plasticity (in ctp8 and further) appeared to stabilize and so did other parameters. The entire process is very similar to the development of animals and human beings.

**Figure 1:**
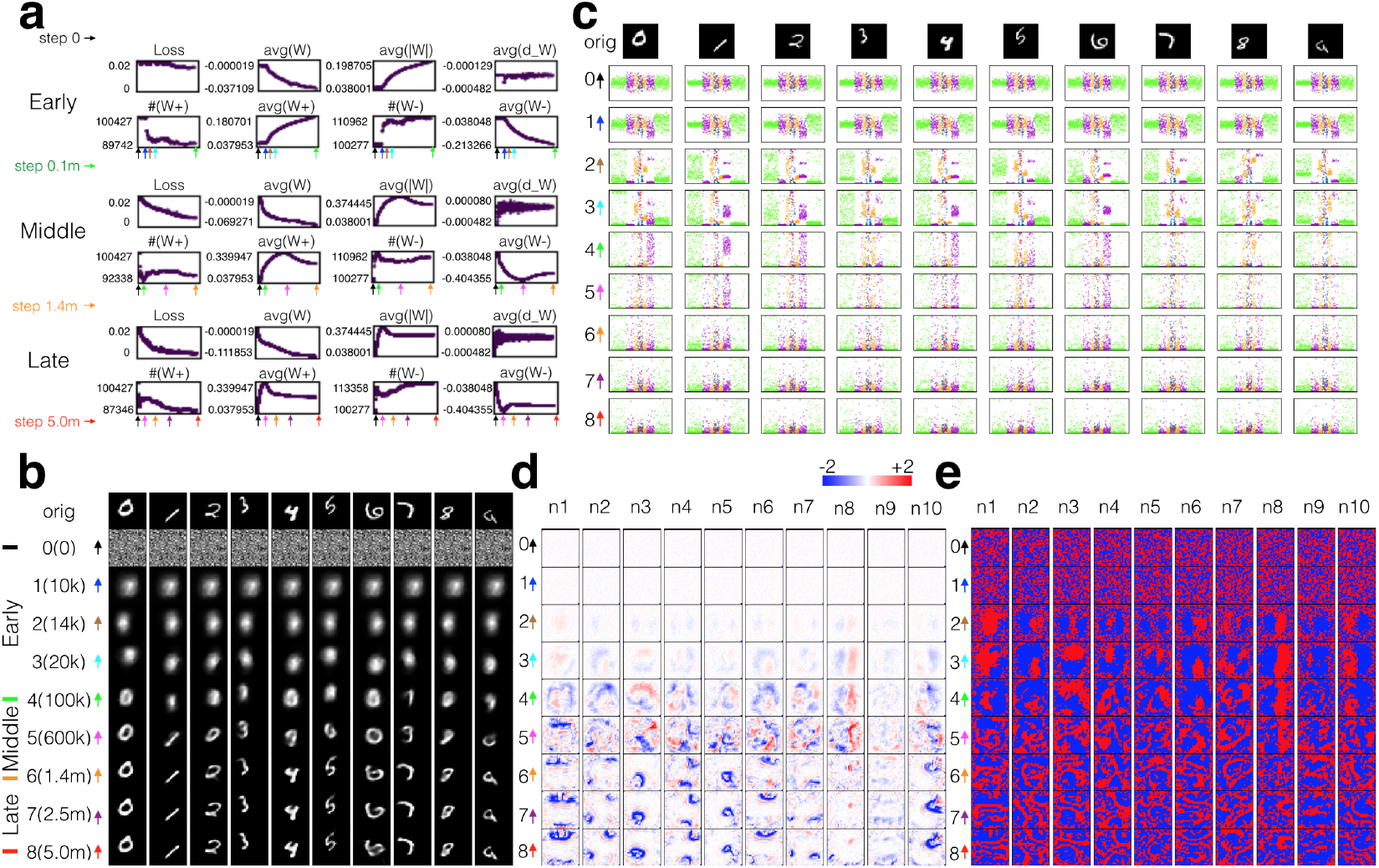
RaI development process. (a) Network parameters evolution over the whole development process (Supplementary Table 1). It contains three stages (early, middle and late stage), and 9 critical time points (ctp) according to the key changes in synaptic weights, neuronal firing patterns, and reconstruction performance. Note, x-axis represents training steps, k for thousand and m for million steps. (b) Digit reconstructions across the development process, where RaI first figured out the digit’s salience map with correct retinal location, size, and orientation in the early stage, then engraved the basic structure in the middle stage, finally refined the structure in a longer duration in the late stage. (c) Evolution of neuronal responses for 10 different digit instances, ranging from initial non-specific firing to final object-related firing. Neurons are presented in order with layers indicated by colors. (d) Evolution of synaptic weights from the retina to the first ten neurons in the e0 layer (Supplementary Figure 2). The increase of weight strength of both excitatory (positive) and inhibitive (negative) synapses facilitated to capture the basic characteristics of digits (ctp0-5), whereas the subsequent decline of weight strength while keeping synaptic pattern sharpen, enabled the recognition of structure details (ctp5-8). (e) Evolution of the distribution of excitatory (red) and inhibitive (blue) synapses (Supplementary Figure 3). It initially shows randomly distributed patterns (ctp0-1), followed by cluster-like patterns (ctp2-4), and finally stripe-like patterns(ctp5-8).

After 5 million steps of training, RaI was mature in recognizing both digit properties and structure details. Then, we looked into the information represented in the Encoding layer to investigate how the digit properties were encoded by these 32 neurons. Figure 2a shows the correlation analysis between retinal position x and neuronal activity across 128 horizontal translation trials. We witness that all layers contain a certain number of x-related (P < 0.01) neurons, which is reasonable and even necessary because precise digit reconstruction requires retinal position information being carried throughout the whole network. 5 out of 32 neurons in the Encoding layer are x-related neurons in Figure 2b-c, in which activity of neuron 24 (n24, the most significantly x related) demonstrates three characteristics: (1) the digit position modulated the activity of n24; (2) the modulation relied on digit identity and morphology; (3) continuous position alteration resulted in continuous change of neuronal response. Therefore, RaI has the diffeomorphic mapping characteristics (no ReLU in RaI) that guarantee its tolerance to the position variation of digit instance. All these are consistent with the property-preserving representation characteristics of inferior temporal (IT) neurons in monkey studies [8, 23].

**Figure 2:**
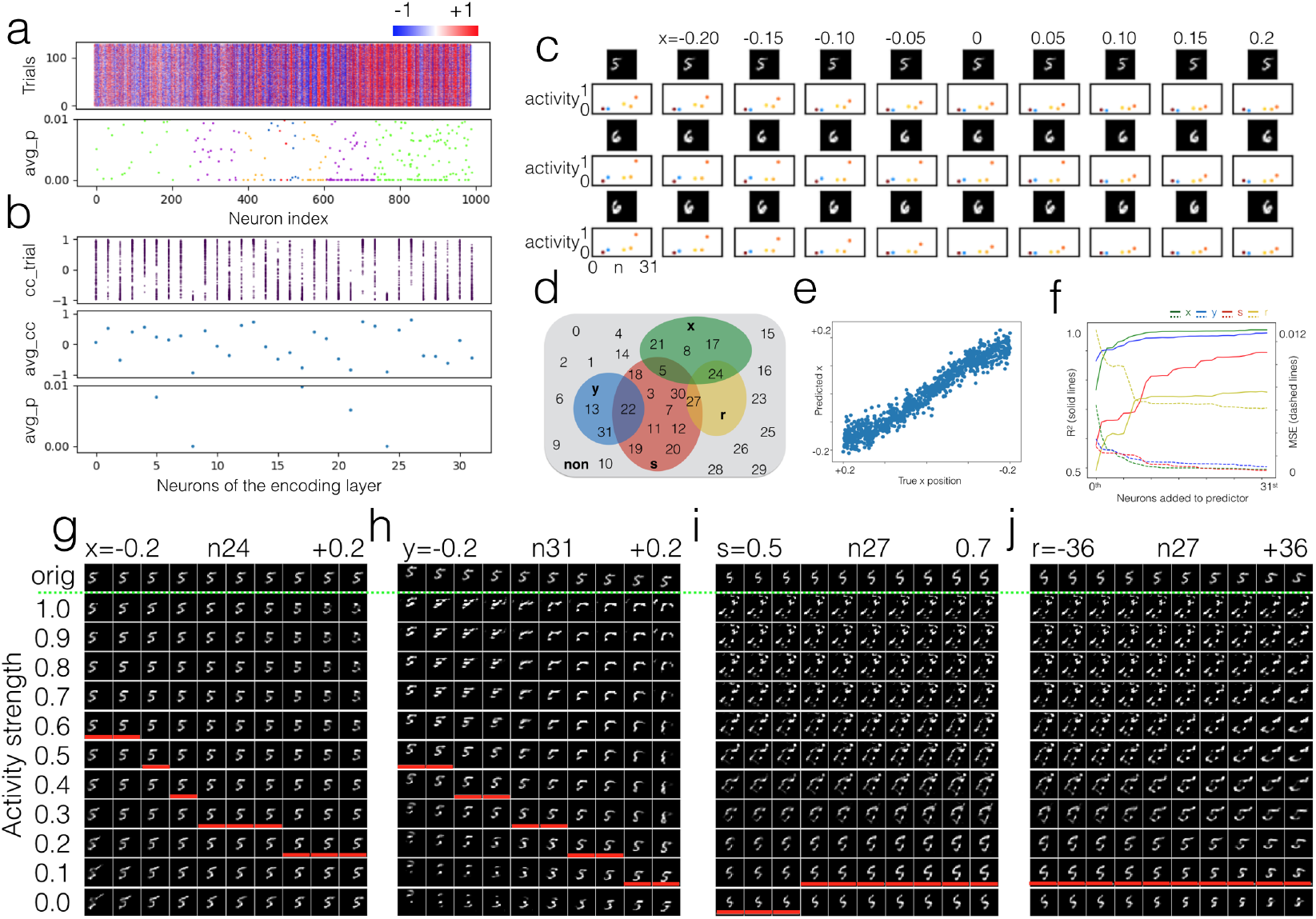
Neuronal responses to digit translation, scaling, and rotation. (a) Neuronal responses in the horizontal translation experiment (top), and x correlated neurons (bottom). (b) The correlation coefficient between neuronal activity and x position for 128 trials (top), where each point represents one trial, and all 32 neurons in the Encoding layer are presented. The middle and bottom panels show the averaged cc across 128 trials and 5 significant (P < 0.01) x-neurons, respectively. (c) x translation, and corresponding responses of x-neurons. Note, activities of neuron 24 (n24, the most significantly x-related) were strongly modulated by digits’ x positions, identities, and even morphologies. (d) Neuronal categorization according to digits’ properties. The non-specific and horizontal translation (x), vertical translation (y), scaling (s), rotation (r) specific neurons took a proportion of 46.9%, 15.6%, 9.4%, 34.4%, 6.3% respectively with certain overlaps. (e) Prediction of position x. The encoding layer contained untangled x information that was easily extracted by a simple linear decoder implemented by a downstream layer. Similar studies were performed for y, s, r in the Supplementary Figure 4-7. (f) Prediction performance (coefficient of determination R2 and mean square error). The retinal position (x, y) were predicted with relatively high accuracy, while scaling and orientation (r, s) with relatively low accuracy. (g) The influence of n24 on digit reconstructions regarding x position. From left column to right, x position increases from −0.2 to +0.2. Images on each row are reconstructed with the modified activities of n24. Red bars highlight the optimal activities derived from RaI. (h-j) similar plots for y, s, and r.

Similar correlations were performed on properties of vertical translation (y), scaling (s), and rotation (r) (Supplementary Figure 4-7), and the 32 neurons were functionally categorized into 5 classes (Figure 2d), which demonstrates the population coding nature of these neurons [24]. To test whether these population responses represent the digit properties, a simple linear summation decoder (Supplementary Figure 1) was implemented by the downstream circuit to predict position x from population responses of the Encoding neurons. Figure 2e demonstrates that the downstream layer could perceive (or predict) the objective x value with a simple linear circuit. Figure 2f shows the predictive performance in terms of R square and mean square error (MSE), demonstrating that perceptions of retinal position x and y were the most accurate, while that of digit size was worse, and in-plane orientation was the worst. Interestingly, although we didn’t train RaI with any digit properties, it could automatically disentangle the manifolds of these properties.

To investigate what information individual neuronal response represents, we modulated the activity of every single neuron, and traced the alteration of image reconstruction. Figure 2g demonstrates how the neuron n24 influences the reconstruction of digit 5 in various x positions, where we have three observations: (1) For each column, even if n24 activity was significantly changed, the reconstructions of digit 5 were always placed in correct x positions, which suggests that activities of neighboring neurons also contained the position information, and n24 activities alone could not influence the reconstruction position. (2) For each column, different n24 activities did substantially change the reconstructed morphology. However, without communicating with neighboring neurons in the same layer, n24 could readily obtain accurate activity from the feedforward inputs of the upstream layer, which suggests the response of n24 represents some features of objects in specific retinal regions. (3) Across Figure 2g, there was no sudden change occurring with regard to the slight movement of digits or changes of n24 activity, which suggests the RaI reconstruction was noise tolerable to digit translation and neuronal activity. Quite similar was the influence of n31 activity on the digit’s y position, but the reconstruction quality of morphology was more sensitive to the variation of n31 activity. Figure 2i-j show that both scaling and rotation were significantly correlated to n27 activity, which maintained fairly low but extremely important activity magnitude around 0.1, and disturbing such magnitude would substantially worsen the reconstruction quality, which suggests both big and small responses are necessary parts of the population coding [25]. In sum, in the population coding scheme, what exactly a single neuronal activity represents can hardly be described, because the interpretation may dramatically alter when neighboring neurons change firing patterns.

To tackle whether neuronal activities encode digit identity, correlation coefficients (cc) between neuronal activities and digit identities were computed (Figure 3a), where nearly all neurons were involved in digit identity coding. The supplementary Figure 8 shows that though variations existed in digit morphologies, these corresponding digit-related neurons had a conservative response pattern regarding each identity. Nevertheless, no single ‘identity’ neuron was found in the Encoding layer. For instance, n14 (the most 7-correlated neuron) showed diverse influence on the reconstruction of digit 7 instances in different conditions (Figure 3b). Then, we used a simple logistic regression layer (Supplementary Figure 1) to assess whether neurons ensemble contains explicit digit identity information. As compared with the baseline accuracy of 10%, this simple logistic regression layer could predict digit with accuracy of 60% (Figure 3c), though no labels were provided during the training process, which indicates that the Encoding neurons are capable of disentangling digit identity which allows downstream layers to easily associate category to specific digit instances. In addition, we found that if the digits were placed in the retinal center, certain rotating or scaling would not drastically deteriorate the recognition accuracy, but placing digits to peripheral regions, especially on the top or bottom of the retina would significantly deteriorate the digit recognition, that is why eye movement is necessary for animal neural network.

**Figure 3:**
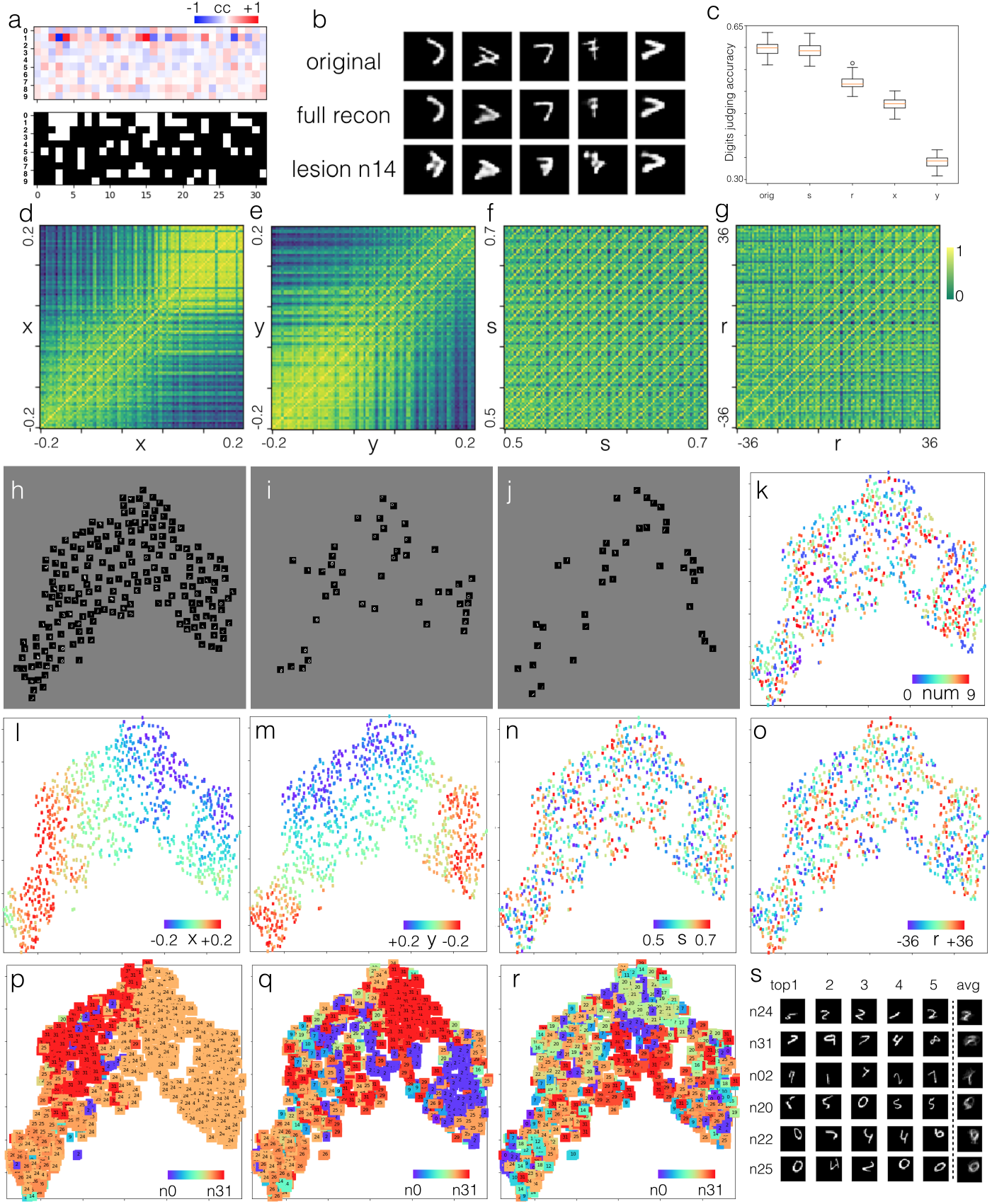
Digit identity recognition. (a) The cc matrix between neuronal activity and digit identity (top), and identity-related (P < 0.01) neurons highlighted in white (bottom). (b) Lesion of n14 (the highest digit7-correlated neuron) damaged the reconstruction of various 7 instances in different degrees. (c) Digits classification accuracy without or with perturbation of x, y, s, r properties. Though training was conducted with no digit labels, a simple logistic classifier could be used to predict digit identity from population responses of the Encoding layer. Note, the accuracy of the natural guess is 10%. (d) Population similarity matrices for x translation. The 100 by 100 cc matrix was computed between population responses across 32 Encoding neurons to 100 stimuli, which were constituted by ten x-position blocks, each contained ten digits from 0 to 9 (supplementary method). Note that the paradiagonal stripes indicate property-invariant digit selectivity. (e-f) Similar matrices for properties of y, s, and r. (h) Non-overlapping view of digit instances in 2D tSNE space. It clearly demonstrates that these images were primarily arranged according to retinal positions, rather than digit identities. (i-j), Distribution of digit 0 and 1 in 2D tSNE space, respectively. (k-o), Distribution of image identities and properties of x, y, s, and r in 2D tSNE space. Retinal position x and y explicitly dominated the global distribution of images, while digit identity, scaling, and orientation showed locally clustering characteristics. (p-r), The first 3 highest responsive neurons for each image in 2D tSNE space. n24 and n31 fired strongest for most digit images, which seemed to encode features in specific retinal regions, while other neurons acted to add or subtract structural details in these regions. (s) Top5 favorite and average (of top20) images. These neurons fired in favor of some features in specific regions, not necessarily related to specific digit identity.

Visual processing needs to handle identity-preserving image transformation problem or to tolerate property variations during identity recognition. To investigate this, we introduced similarity matrices [6] between population responses with gradually changed properties (supplementary method). One remarkable feature in Figure 3d is the presence of paradiagonal stripes, which indicates the same digit with decent horizontal displacement will not severely disturb the population response pattern. The other feature is the high correlation region on the top right of Figure 3d, which indicates that digits placed on the right part of retina will elicit similar neuronal activities regardless of their identities. Vertical variation of digit position has resembled similarity matrix in Figure 2e. Nevertheless, the variation of digit size demonstrates remarkably more and stronger paradiagonal stripes than others, which suggests that same digit instances but different sizes were encoded by similar population response patterns, and only slight modulation on population activities was needed for recognizing the size alteration. In contrast, relatively large adjustment of a coordinated population response was needed to recognize retinal position changes, which is consistent with supplementary Figure 9. So the natural neural mechanism in handling the translational invariance problem is not based on max-pooling but on coordinated adjustment of population responses.

The t-Distributed Stochastic Neighbor Embedding (tSNE) could provide visible insights into how these digit instances organized in a 2D state space [26]. Figure 3h-j illustrate two main findings: (1) retinal position x and y dominated the global distribution of digit in the tSNE space; (2) digit identities had a tendency to cluster locally. So alteration of the retinal position indicated a big move in tSNE space, which required a large scale adjustment of coordinated responses to achieve it; while alteration of digit size meant a small nudge in the tSNE space, which required little adjustment in population responses and showed remarkably more and stronger paradiagonal strips. From Figure 3p-r, we witness that activities of n24, n31, n2 and n25 might provide location guidance information for image reconstruction, while other neurons provided relatively small activities to refine the structure in such regions. However, both big and small responses were integral parts of population coding. Figure 3s (Supplementary Figure 10) shows that neurons of the Encoding layer fire in favor of features in a specific retinal region rather than digit identities.

Learning new and forgetting old skills (or knowledge) are basic characteristics of animal brains. Here, we will reveal how new recognition skills are added into the working network while previous skills are still preserved without big disruptions. It is not surprising that, in the beginning, RaI would reflectively treat novel structures as some known digits. In Figure 4a, RaI treated the square as compacted digit 8 and double-digit as a wide single digit. Figure 4b demonstrates that novel structures (symbol ‘x’) induced significantly (P = 2.43e-50) stronger (2.5 times) synaptic plasticity than known structures (digits). More importantly, these changes occurred on all synapses in relatively small magnitude, so that previous sharp synaptic patterns could be retained. This guarantees that those known structures could still elicit similar neuronal responses, which enable downstream layers to process data properly and robustly without big modulation. This also guarantees that any novel structure is learned and recognized by the population coding scheme.

**Figure 4:**
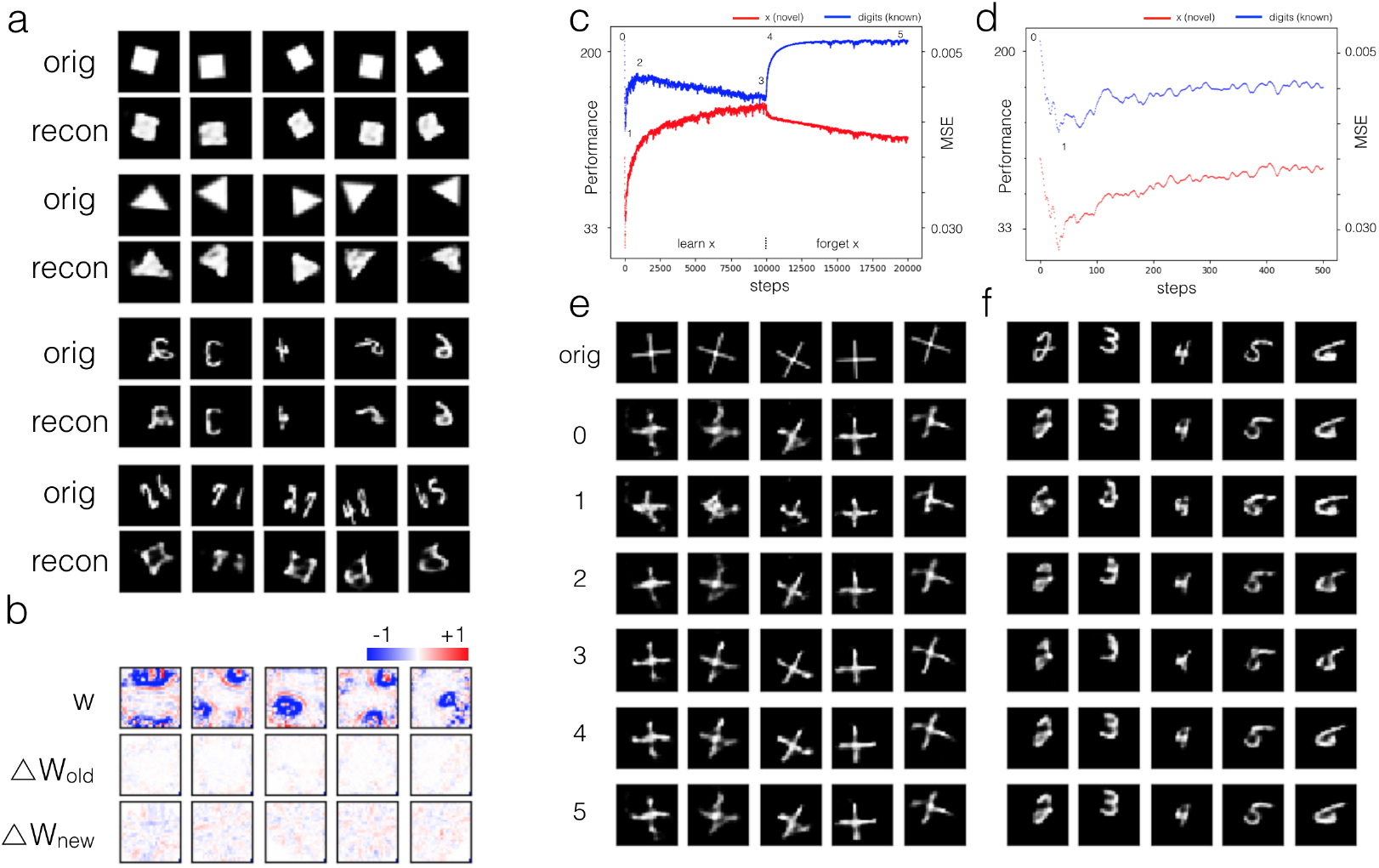
Learning and forgetting. (a) Reconstruction of new structures including solid square, triangle, mirrored digits, and double digits. (b) Synaptic weights distribution and changes after 10,000 steps training of known and novel (symbol ‘x’) structures. (c) Performance evolution during training and forgetting processes. RaI was trained with novel symbol x for the first 10,000 steps, followed by known digits in the next 10,000 steps. Red represents test performance (or MSE in the upside-down y-axis) of novel structure x, while blue represents known digits. Six distinguished ctps (0-5) were observed across the entire process. (d) Enlarged view of the first 500 steps which clearly demonstrates the initial drop and subsequent rise of the reconstruction performance. (e-f) The reconstruction of symbol x and digits for 6 ctps, which demonstrates the performance evolution.

The connection rewiring process during learning and forgetting involved several key stages (Figure 4c-f). At ctp0, RaI could recognize and reconstruct both novel and known structures with relatively good quality. Surprisingly, the first 30 steps training of symbol ‘x’ induced a sudden drop of performance for both structures. The novel experience made RaI unstable until ctp1 when RaI found a new attractor to settle on [2], thenceforth a new round of connectivity adjustment started. In transition 1-2, RaI rapidly improved the performance for symbol x because symbol x instances were the only input to the network. Unexpectedly, the performance for known digits also got improved, which might be due to the synaptic adjustment for better reuse of previous connectivity. After sufficient recovery of RaI capacity, performance for known structure began to decline again until ctp3. After that, RaI began to experience only digit instances again, the network soon recovered the digit recognition performance to the original level, while the forgetting of symbol x took effect rapidly. For transition 4-5 and further, the input of digits continued to enhance the digit recognition while decrease that of symbol x. Note that the real neural network is much larger, the performance alteration (blue curve from ctp0 to ctp3) would not change such dramatically. Here, we want to differentiate between recognition learning and knowledge learning, where the former (like tasting spicy will gradually lower the perception of spicy taste) is a long term process requiring plasticity of sensory synapses along processing stream, while the latter (like learning a new foreign word) could be a short process requiring certain new association connectivity formed between sensory and downstream regions.

## 3 Conclusion

In this paper, we proposed a recognition and imagination network as an analogous model to study the animal-like visual mechanism. It is the first time that the responsive characteristics of an artificial network matched the electrophysiological measurements in monkeys, and the first time to interpret how such animal-like neuronal responses code objects. In the network development study, RaI experienced distinct developmental stages, characterized by specificities in neural types, connectivity strength, firing patterns, from an early stage of coarse salience map recognition to mature stage of fine structure recognition. RaI automatically developed animal-like neural mechanisms such as sparse coding, synaptic prune, localized receptive field, and population coding schemes, etc. In the object recognition study, we found that RaI used a population coding scheme, where each unit responded concertedly to features in a specific retinal region, which allowed to robustly untangle the input images and to form an object invariance representation. And from the population responses, object identity and properties could be accurately extracted by a simple linear or logistic regressor implemented by a downstream circuit. In the learning and forgetting study, novel structures induced plasticity over entire synapses in low magnitude while kept original specificity of synaptic weights unchanged, which guaranteed that new knowledge learning would not disrupt the old ones. In sum, Under the same principle of ‘free energy minimization’, animal and RaI could independently evolve similar neural networks, with matched observations in developments, neuronal response, and learning characteristics, etc. This work benefits the understanding of the human neural mechanism and the development of human-like intelligence.

## Notes

#### Summary of Updates

UPDATE A NEW VERSION

